# Genomics-enabled analysis of the emergent disease cotton bacterial blight

**DOI:** 10.1101/127670

**Authors:** Anne Z. Phillips, Jeffrey C. Berry, Mark C. Wilson, Anupama Vijayaraghavan, Jillian Burke, J. Imani Bunn, Tom W. Allen, Terry Wheeler, Rebecca Bart

**Author notes:** these authors contributed equally to this work (Jeffrey C. Berry, Mark C. Wilson).

## Abstract

Cotton bacterial blight (CBB), an important disease of (*Gossypium hirsutum*) in the early 20^th^ century, had been controlled by resistant germplasm for over half a century. Recently, CBB re-emerged as an agronomic problem in the United States. Here, we report analysis of cotton variety planting statistics that indicate a steady increase in the percentage of susceptible cotton varieties grown each year since 2009. Phylogenetic analysis revealed that strains from the current outbreak cluster with race 18 *Xanthomonas citri* pv. *malvacearum* (*Xcm*) strains.

Illumina based draft genomes were generated for thirteen *Xcm* isolates and analyzed along with 4 previously published *Xcm* genomes. These genomes encode 24 conserved and nine variable type three effectors. Strains in the race 18 clade contain 3 to 5 more effectors than other *Xcm* strains. SMRT sequencing of two geographically and temporally diverse strains of *Xcm* yielded circular chromosomes and accompanying plasmids. These genomes encode eight and thirteen distinct transcription activator-like effector genes. RNA-sequencing revealed 52 genes induced within two cotton cultivars by both tested *Xcm* strains. This gene list includes a homeologous pair of genes, with homology to the known susceptibility gene, MLO. In contrast, the two strains of *Xcm* induce different clade III SWEET sugar transporters. Subsequent genome wide analysis revealed patterns in the overall expression of homeologous gene pairs in cotton after inoculation by *Xcm*. These data reveal host-pathogen specificity at the genetic level and strategies for future development of resistant cultivars.

**Author Summary:** Cotton bacterial blight (CBB), caused by *Xanthomonas citri* pv. *malvacearum* (*Xcm*), significantly limited cotton yields in the early 20^th^ century but has been controlled by classical resistance genes for more than 50 years. In 2011, the pathogen re-emerged with a vengeance. In this study, we compare diverse pathogen isolates and cotton varieties to further understand the virulence mechanisms employed by *Xcm* and to identify promising resistance strategies. We generate fully contiguous genome assemblies for two diverse *Xcm* strains and identify pathogen proteins used to modulate host transcription and promote susceptibility. RNA-Sequencing of infected cotton reveals novel putative gene targets for the development of durable *Xcm* resistance. Together, the data presented reveal contributing factors for CBB re-emergence in the U.S. and highlight several promising routes towards the development of durable resistance including classical resistance genes and potential manipulation of susceptibility targets.

## Introduction

Upland cotton (*Gossypium hirsutum* L.) is the world’s leading natural fiber crop. Cotton is commercially grown in over 84 countries, and in the United States, is responsible for $74 billion annually [1, 2]. Numerous foliar diseases affect cotton throughout the world’s cotton growing regions. Historically, one of the most significant foliar diseases has been bacterial blight, caused by *Xanthomonas citri* pv. *malvacearum*. Cotton bacterial blight significantly limited cotton yield in the late 20^th^ century. In the 1940’s and 1950’s, breeders identified and introgressed multiple resistance loci into elite germplasm [3-5]. This strategy proved durable for over half a century. In 2011, cotton bacterial blight (CBB) returned and caused significant losses to farmers in the southern United States, including in Arkansas and Mississippi. Nonetheless, CBB has received little research focus during the last several decades because, prior to 2011, losses from this disease were not substantial. Modern molecular and genomic technologies can now be employed expeditiously to deduce the underlying cause of the disease re-emergence and pinpoint optimized routes towards the development of durable resistance.

CBB is caused by *X. citri pv. malvacearum* (*Xcm*); however, the pathogen has previously been placed within other species groupings [6-9]. The *Xcm* pathovar can be further divided into at least 19 races according to virulence phenotypes on a panel of historical cotton cultivars: Acala-44, Stoneville 2B-S9, Stoneville 20, Mebane B-1, 1-10B, 20-3, and 101-102.B [10, 11]. Historically, the most common race observed in the U.S. has been race 18, which was first isolated in 1973 [12]. This race is highly virulent, causing disease on all cultivars in the panel except for 101-102.B. However, this diagnostic panel of cotton varieties used to race type strains is no longer available from the USDA/ARS, Germplasm Resources Information Network (GRIN).

CBB can occur at any stage in the plant’s life cycle and on any aerial organ. Typical symptoms include seedling blight as either pre- or post-emergent damping-off, black arm on petioles and stems, water-soaked spots on leaves and bracts, and most importantly boll rot [10]. The most commonly observed symptoms are the angular-shaped lesions on leaves that can coalesce and result in a systemic infection. Disease at each of these stages can cause yield losses either by injury to the plant or direct damage to the boll. No effective chemical treatments for the disease have been released to date. Methods to reduce yield loss as a result of CBB include acid de-linting cotton seed prior to planting, field cultivation practices to reduce sources of overwintering inoculum and planting cultivars with known sources of resistance [3, 4, 8, 13, 14].

Xanthomonads assemble the type three secretion system (T3SS), a needle-like structure, to inject diverse type three effectors (T3Es) into the plant cell to suppress immunity and promote disease [15-19]. For example, transcription activator-like (TAL) effectors influence the expression levels of host genes by binding directly to promoters in a sequence-specific way [20]. Up-regulated host genes that contribute to pathogen virulence are termed susceptibility genes and may be modified through genome editing for the development of resistant crop varieties [21].

Plants have specialized immune receptors, collectively known as nucleotide-binding leucine rich repeat receptors that recognize, either directly or indirectly, the pathogen effector molecules [22, 23]. Historically, this host-pathogen interaction has been termed the ‘gene-for-gene’ model of immunity, wherein a single gene from the host and a single gene from the pathogen are responsible for recognition [24]. Recognition triggers a strong immune response that often includes a localized hypersensitive response (HR) in which programmed cell death occurs around the infection site [25]. Nineteen CBB resistance loci have been reported in *Gossypium hirsutum* breeding programs; however, none have been molecularly identified [8, 13].

Here we combine comparative genomics of the pathogen *Xcm* with transcriptomics of the host to identify molecular determinants of Cotton Bacterial Blight. This will inform the development of durable resistance strategies.

## Results

### CBB Reemergence in the US

In 2011, farmers, extension specialists, and certified crop advisers in Missouri, Mississippi, and Arkansas observed cotton plants exhibiting symptoms of CBB. Widespread infected plant material was observed throughout much of the production area, but appeared to be centered around Clarksdale, Mississippi. In figure 1, we collate reports from this outbreak and overlay these data with US cotton planting statistics to reveal that this disease has spread through much of the cotton belt in the southern U.S. (Figs 1 and S1, Table S1). To date, CBB has been reported from at least eight out of the sixteen states that grow cotton (Fig 1). In 2014, we collected diseased cotton leaves from two sites across Mississippi and proved pathogen causality following Koch’s postulates [26]. In addition, PCR amplification of the 16S rRNA gene confirmed that the causal agent was a member of the *Xanthomonas* genus. Multi locus sequence type (MLST) analysis and maximum-likelihood analysis were performed using concatenated sections of the *gltA*, *lepA*, *lacF*, *gyrB*, *fusA* and *gap-1* loci for increased phylogenetic resolution (Fig 2a). The newly sequenced strains were named MS14002 and MS14003 and were compared to four previously published *Xcm* genomes and thirty-six additional *Xanthomonas* genomes representing thirteen species (Tables 1, S2). MS14002 and MS14003 grouped with the previously published *Xcm* strains as a single unresolved clade, further confirming that the current disease outbreak is CBB and is caused by *Xcm*. The species designation reported here is consistent with previous reports [6, 7].

**Figure 1.**
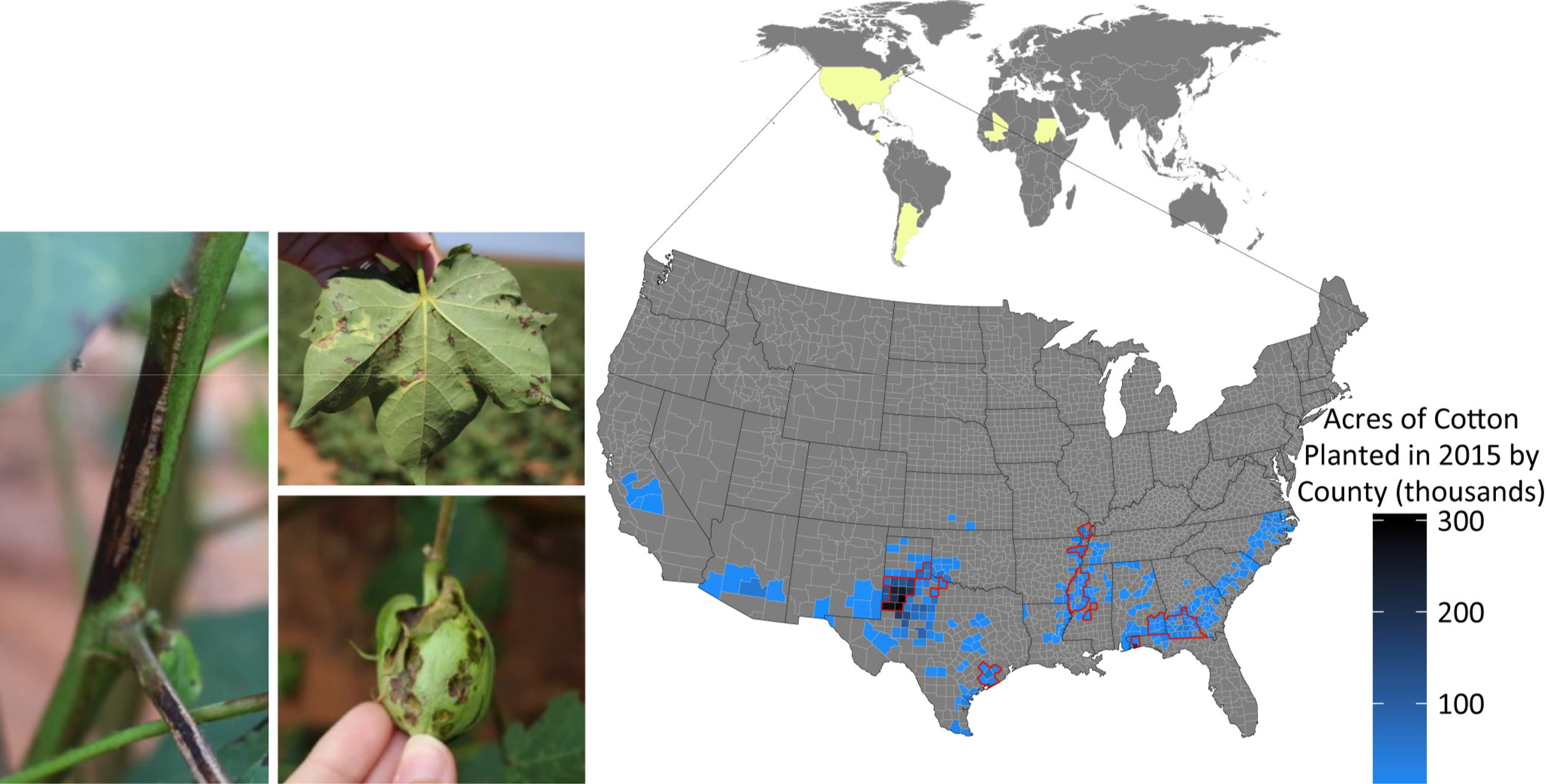
Cotton Bacterial Blight (CBB) symptoms and reemergence across the southern United States. (Left) Typical CBB symptoms present in cotton fields near Lubbock, TX during the 2015 growing season include angular leaf spots, boll rot, and black arm rot. Yellow shading within world map (top) indicates origin of strains included in this study. Acres of cotton planted per county in the United States in 2015 (blue) and counties with confirmed CBB in 2015 (red outline). Statistics on the area of cotton planted in the U.S. were acquired from the USDA. CBB was reported by extension agents, extension specialists, and certified crop advisers in their respective states.

**Table 1.** Illumina and SMRT sequenced Xcm genomes described in this paper.

| Strain Name | Identifier | Lat. | Long. | City/Country | Year | Provider | Collector | Platform | Contig # | Avg Contig Len | Total Bases | n50 |
| --- | --- | --- | --- | --- | --- | --- | --- | --- | --- | --- | --- | --- |
| MS14002 |  | 34.12 | -90.52 | Clarksdale, MS | 2014 | R.Bart | R.Bart | Illumina | 545 | 9443.27 | 5146580 | 62542 |
| MS14003 |  | 32.95 | -90.51 | Wilzone, MS | 2014 | R.Bart | R.Bart | Illumina | 2577 | 1511.35 | 3894744 | 2209 |
| Race1 |  |  |  | US |  | S.lu | P.Thaxton | Illumina | 523 | 10127.35 | 5296606 | 48599 |
| Race2 |  |  |  | US |  | S.lu | P.Thaxton | Illumina | 387 | 13402.57 | 5186796 | 54804 |
| Race3 |  |  |  | US |  | S.lu | P.Thaxton | Illumina | 725 | 7207.34 | 5225324 | 28344 |
| Race12 |  |  |  | US |  | S.lu | P.Thaxton | Illumina | 632 | 8134.35 | 5140911 | 21428 |
| Race18 |  |  |  | US |  | S.lu | P.Thaxton | Illumina | 369 | 13924.03 | 5137968 | 112543 |
| AR81009 | CFBP2035 |  |  | Argentina | 1981 | CIRM-CFBP | Frossard P | Illumina | 306 | 17182.59 | 5257872 | 86594 |
| MA81010 | CFBP2036 |  |  | Mali | 1981 | CIRM-CFBP | Frossard P | Illumina | 584 | 9033.09 | 5275326 | 23323 |
| SU58011 | CFBP2530 |  |  | Sudan | 1958 | CIRM-CFBP | Last F.T. M2519 | Illumina | 1134 | 4563.33 | 5174819 | 9682 |
| SU9012 | CFBP5637 |  |  | Sudan | 1992 | CIRM-CFBP | Schmit J. 11781 | Illumina | 377 | 13919.54 | 5247665 | 88522 |
| Xcm013 | MSCT4 |  |  | US |  | S.lu |  | Illumina | 2169 | 2151.5 | 4666607 | 3869 |
| Xcm014 | MSCT8 |  |  | US |  | S.lu |  | Illumina | 580 | 8929.58 | 5179156 | 88255 |
| MS14003 |  | 32.95 | -90.51 | Wilzone, MS | 2014 | R.Bart | R.Bart | SMRT | 4 | 1286176.5 | 5144706 | 5029617 |
| AR81009 | CFBP2035 |  |  | Argentina | 1981 | CIRM-CFBP | Frossard P | SMRT | 4 | 1352212 | 5408848 | 5267057 |

### Contemporary U.S. *Xcm* strains cluster phylogenetically with historical race 18 strains

Race groups have been described for *Xcm* strains by analyzing compatible (susceptible) and incompatible (resistant) interactions on a panel of seven cotton cultivars. Different geographies often harbor different pathogen races [7]. Consequently, one possible explanation for the recent outbreak of CBB would be the introduction of a new race of *Xcm* capable of overcoming existing genetic resistance. Only 2 varieties of the original cotton panel plus three related cultivars, were available and these cultivars were not sufficient to determine whether a new race had established within the U.S. Thirteen *Xcm* strains were sequenced using Illumina technology to determine the phylogenetic relationship between recent isolates of *Xcm* and historical isolates. Isolates designated as race 1, race 2, race 3, race 12 and race 18 have been maintained at Mississippi State University with these designations. Additional isolates were obtained from the Collection Française de Bactéries associées aux Plantes (CFBP) culture collection. Together, these isolates include nine strains from the US, three from Africa, and one from South America and span collection dates ranging from 1958 through 2014 (Fig 1, Table 1). Illumina reads were mapped to the *Xanthomonas citri* subsp. *citri* strain Aw12879 (Genbank assembly accession: GCA_000349225.1) using Bowtie2 and single nucleotide polymorphisms (SNPs) were identified using Samtools [27, 28]. Only regions of the genome with at least 10x coverage for all genomes were considered. This approach identified 17,853 sites that were polymorphic in at least one genome. Nucleotides were concatenated and used to build a neighbor-joining tree (Fig 2b). This analysis revealed that recent U.S. *Xcm* isolates grouped with the race 18 clade. Notably, the race 18 clade is phylogenetically distant from the other *Xcm* isolates.

**Figure 2.**
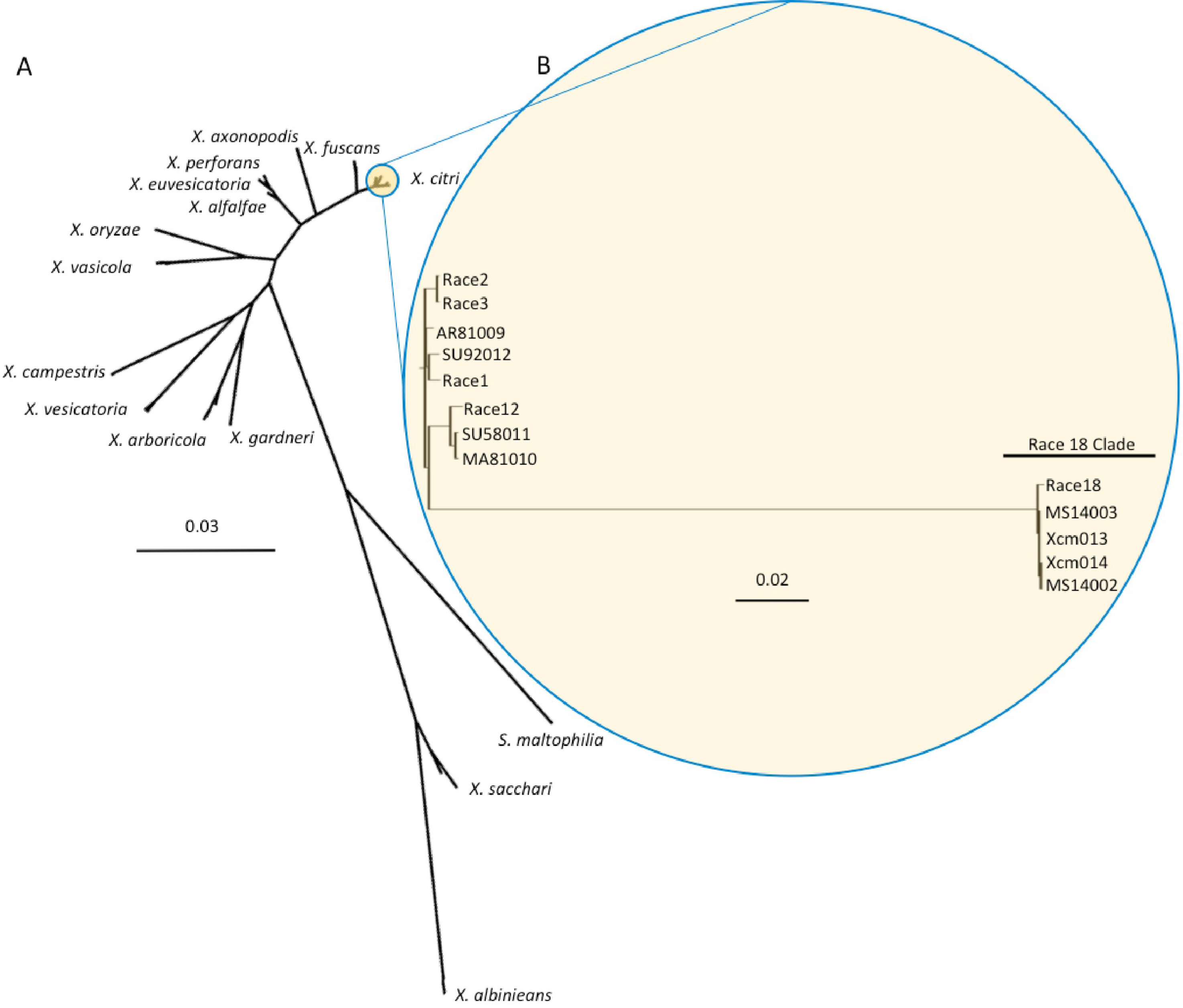
Phylogenetic analysis of *Xcm* isolates and 13 species of *Xanthomonas*. A) MLST (Multi Locus Sequence Typing) and maximum likelihood analysis of 13 Illumina sequenced *Xcm* isolates (this paper) and 40 other Xanthomonads using concatenated sections of the gltA, lepA, lacF, gyrB, fusA and gap-1 loci. B) SNP based neighbor-joining tree generated from 17,853 variable loci between 13 *Xcm* isolates and the reference genome *Xanthomonas citri* subsp. *citri* strain Aw12879. The tree was made using the Simple Phylogeny tool from ClustalW2.

### Contemporary US Xcm strains have conserved type three virulence protein arsenals and disease phenotypes with historical race 18 strains

Xanthomonads deploy many classes of virulence factors to promote disease. Type three effectors (T3E) are of particular interest for their role in determining race designations. T3E profiles from sixteen *Xcm* isolates were compared to determine whether a change in the virulence protein arsenal of the newly isolated strains could explain the re-emergence of CBB. Genomes from 13 *Xcm* isolates were *de novo* assembled with SPAdes and annotated with Prokka based on annotations from the *X. euvesicatoria* (aka. *X. campestris* pv. *vesicatoria*) 85-10 genome (NCBI accession: NC_007508.1). T3Es pose a particular challenge for reference based annotation as no bacterial genome contains all effectors. Consequently, an additional protein file containing known T3Es from our previous work was included within the Prokka annotation pipeline [15, 29]. This analysis revealed 24 conserved and 9 variable *Xcm* T3Es (Fig 3a). Race 18 clade isolates contain more effectors than other isolates that were sequenced. The recent *Xcm* isolates (MS14002 and MS14003) were not distinguishable from the historical race 18 isolate, with the exception of XcmNI86 isolated from Nicaragua in 1986, which contains mutations in XopE2 and XopP.

**Figure 3.**
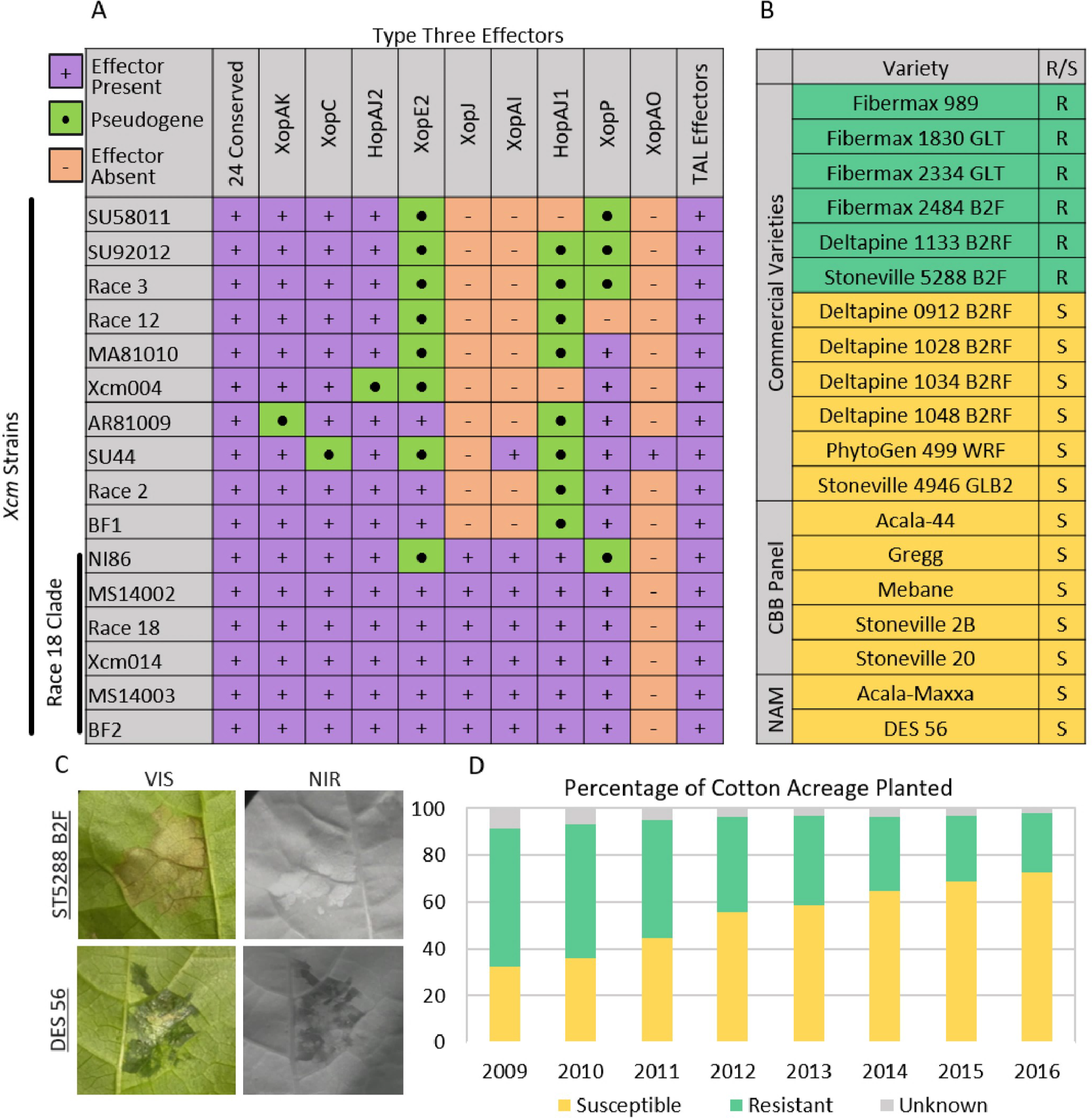
Molecular and phenotypic analysis of *Xcm* and *G. hirsutum* interactions. A) Type three effector profiles of *Xcm* isolates were deduced from *de novo*, Illumina based genome assemblies. Effector presence or absence was determined based on homology to known type three effectors using the program Prokka. B) Commercial and public *G. hirsutum* cultivars were inoculated with 13 *Xcm* isolates. Susceptible (S) indicates water soaking symptoms. Resistant (R) indicates a visible hypersensitive response. Plants were screened with a range of inoculum concentration from OD_600_ = 0.001-0.5. C) Disease symptoms on *G. hirsutum* cultivars Stoneville 5288 B2F and DES 56 after inoculation with *Xcm* strain AR81009 (OD_600_ = 0.05). Symptoms are visualized under visible (VIS) and near infrared (NIR) light. D) The proportion of US fields planted with susceptible and resistant cultivars of *G. hirsutum* was determined using planting acreage statistics from the USDA-AMA and disease phenotypes based on previous reports for common cultivars [34-36].

Analysis of the genomic sequence of T3Es revealed presence/absence differences, frameshifts and premature stop codons. However, this analysis does not preclude potential allelic or expression differences among the virulence proteins that could be contributing factors to the re-emergence of CBB. Therefore, newly isolated strains may harbor subtle genomic changes that have allowed them to overcome existing resistance phenotypes. Many commercial cultivars of cotton are reported to be resistant to CBB [30-32]. Based on these previous reports, we selected commercial cultivars resistant and susceptible (6 of each) to CBB. In addition, we included 5 available varieties that are related to the historical panel as well as 2 parents from a nested association mapping (NAM) population currently under development [33]. All varieties inoculated with the newly isolated *Xcm* strains exhibited inoculation phenotypes consistent with previous reports (Figs 3b,c). In these assays, bright field and near infrared (NIR) imaging were used to distinguish water-soaked disease symptoms from rapid cell death (HR) that is indicative of an immune response. These data confirm that existing resistance genes present within cotton germplasm are able to recognize the newly isolated *Xcm* strains and trigger a hypersensitive response. Together, the phylogenetic analysis, effector profile conservation and cotton inoculation phenotypes, confirm that the recent outbreak of *Xcm* in the US represents a re-emergence of a race 18 clade *Xcm* and is not the result of a dramatic shift in the pathogen.

The USDA Agricultural Marketing Service (AMS) releases reports on the percentage of upland cotton cultivars planted in the U.S. each year (www.ams.usda.gov/mnreports/cnavar.pdf). Most of these varieties are screened for resistance or susceptibility to multiple strains of *Xcm* by extension scientists and published in news bulletins [30, 31, 34-38]. These distinct datasets were cross referenced to reveal that only 25% of the total cotton acreage was planted with resistant cultivars in 2016 (Fig 3d, Table S3). This is part of a larger downward trend in which the acreage of resistant cultivars has fallen each year since at least 2009 when the percentage of acreage planted with resistant varieties was at 75%.

### Comparative genome analysis for two Xcm strains

Differences in virulence were observed among *Xcm* strains at the molecular and phenotypic level. In order to gain insight into these differences, we selected two strains from our collection that differed in T3E content, virulence level, geography of origin and isolation date. AR81009 was isolated in Argentina in 1981 and is one of the most virulent strains investigated in this study; MS14003 was isolated in Mississippi in 2014 and is a representative strain of the race 18 clade (Fig S2). The latter strain causes comparatively slower and diminished leaf symptoms; however, both strains are able to multiply and cause disease on susceptible varieties of cotton (Fig S3). Full genome sequences were generated with Single Molecule Real-Time (SMRT) sequencing. Genomes were assembled using the PacBio Falcon assembler which yielded circular 5Mb genomes and associated plasmids. Genic synteny between the two strains was observed with the exception of two 1.05 Mb inversions (Fig 4). Regions of high and low GC content, indicative of horizontal gene transfer, were identified in both genomes. In particular, a 120kb insertion with low GC content was observed in AR81009. This region contains one T3E as well as two annotated type four secretion system-related genes, two conjugal transfer proteins, and two multi drug resistant genes (Fig 4 insert). MS14003 contains three plasmids (52.4, 47.4, and 15.3kb) while AR81009 contains two plasmids (92.6 and 22.9kb). Analysis of homologous regions among the plasmids was performed using progressiveMauve [39]. This identified four homologous regions greater than 1kb that were shared among multiple plasmids (Fig 4).

**Figure 4.**
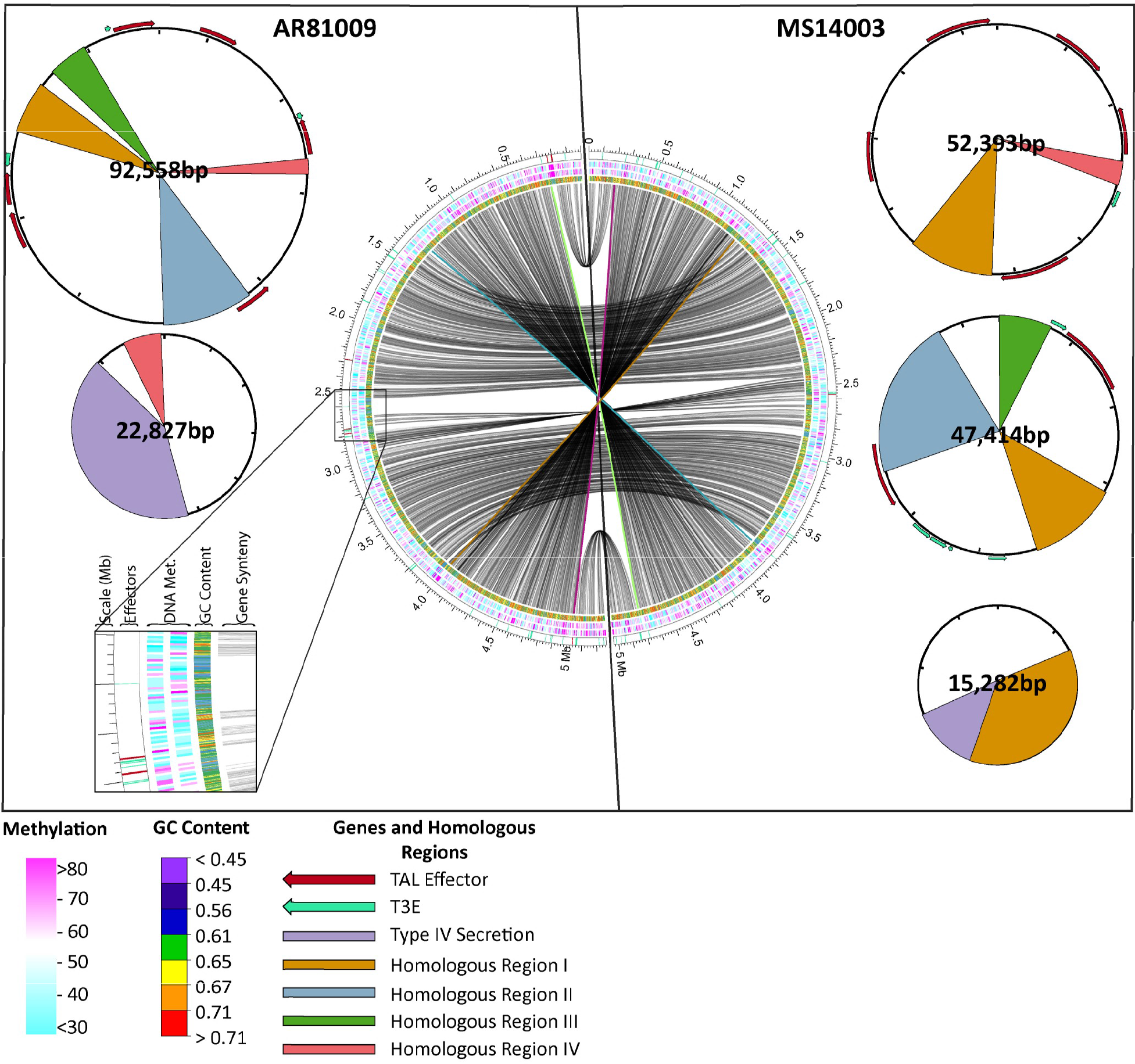
SMRT sequencing of two phenotypically and geographically diverse *Xcm* isolates: MS14003 and AR81009. Circos plot visualization of two circular *Xcm* genomes. Tracks are as follows from inside to outside: synteny of gene models; GC Content; DNA Methylation on + and – strands; location of type three effectors (teal) and TAL effectors (red), and position. On each side, accompanying plasmids are cartooned. Type three effector repertoires and the type IV secretion systems were annotated using Prokka. Homologous regions greater than 1kb were identified using MAUVE, and TAL effectors were annotated using AnnoTALE.

Both strains express TAL effector proteins as demonstrated through western blot analysis using a TAL effector specific polyclonal antibody (Fig 5) [40]. However, the complexity of TAL effector repertoires within these strains prevented complete resolution of each individual TAL effector. The long reads obtained from SMRT sequencing are able to span whole TAL effectors, allowing for full assemblies of the TAL effectors in each strain. The AR81009 genome encodes twelve TAL effectors that range in size from twelve to twenty three repeat lengths, six of which reside on plasmids. The MS14003 genome encodes eight TAL effectors that range in size from fourteen to twenty eight repeat lengths, seven of which reside on plasmids (Fig 5). Three partial TAL effector-like coding sequences were also identified within these genomes and are presumed to be non-functional. A 1-repeat gene with reduced 5’ and 3’ regions was identified in both strains directly upstream of a complete TAL effector. In addition, a large 4kb TAL effector was identified in AR81009 with a 1.5 kb insertion and 10 complete repeat sequences. The tool AnnoTALE was used to annotate and group TAL effectors based on the identities of the repeat variable diresidues (RVDs) in each gene [41]. Little homology was identified among TAL RVD sequences within and between strains; only two TAL effectors were determined to be within the same TAL class between strains (TAL19b of AR81009 and TAL19 of MS14003) and two within strain MS14003 (TAL14b and TAL16).

**Figure 5.**
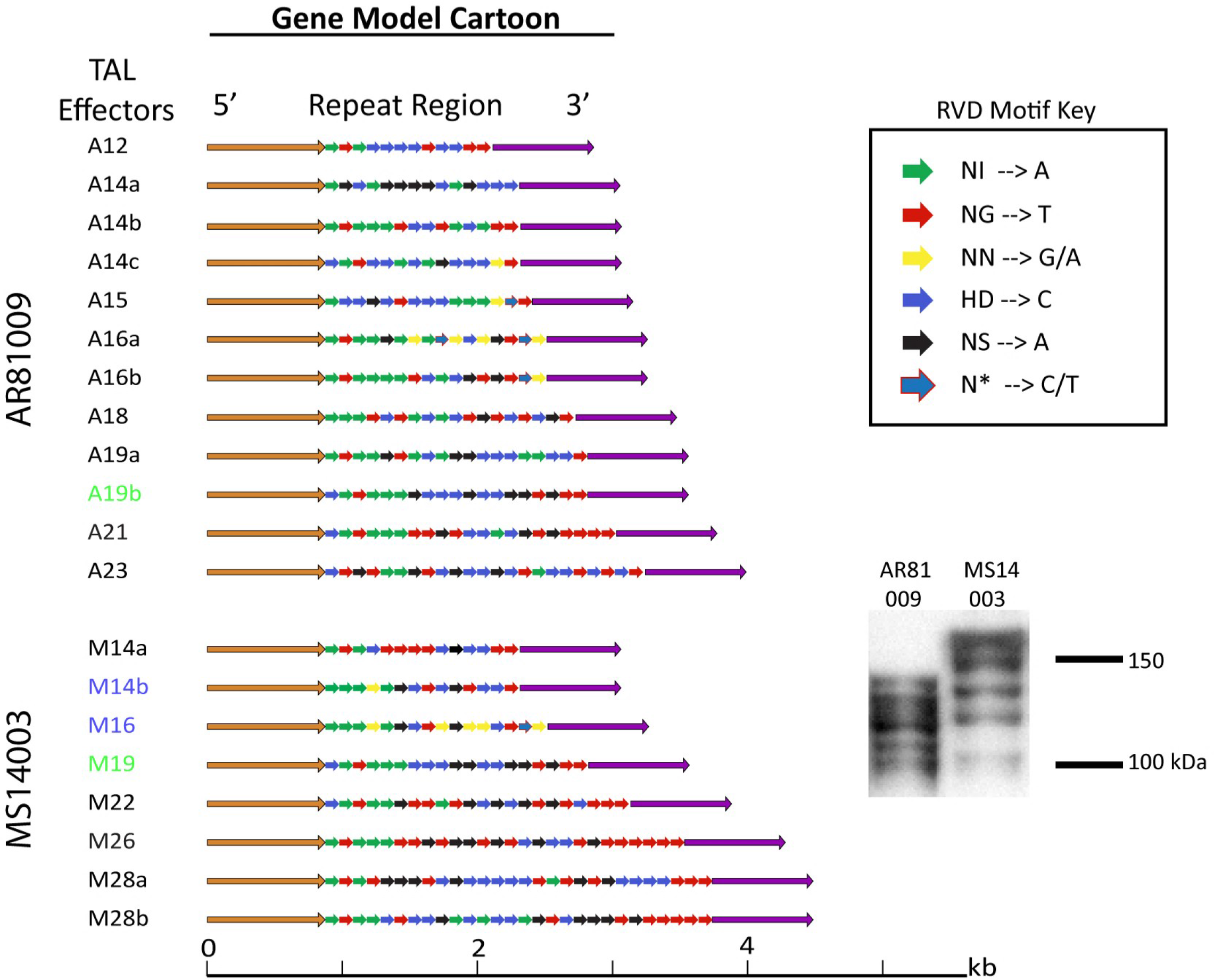
SMRT sequencing and western blot reveal diverse TAL effector repertoires between *Xcm* strains MS14003 and AR81009. Western Blot of TAL effectors using polyclonal TAL-specific antibody and gene models of TAL effectors identified by AnnoTALE. Blue and Green highlighted gene models represent TALs grouped in the same clade by repeat variable di-residue (RVD) sequence using AnnoTALE.

### Transcriptome changes induced by Xcm in G. hirsutum

An RNA-sequencing experiment was designed to determine whether AR81009 and MS14003 incite different host responses during infection (Fig 6a, b). Isolates were inoculated into the phylogenetically diverse *G. hirsutum* cultivars Acala Maxxa and DES 56 [33]. Infected and mock-treated tissue were collected at 24 and 48 hours post inoculation. First, we considered global transcriptome patterns of gene expression. Fifty-two genes were determined to be induced in all *Xcm-G. hirsutum* interactions at 48 hours (Fig 6c, Table S4). Of note among this list is a homeologous pair of genes with homology to the known susceptibility target MLO [42-45]. Gene induction by a single strain was also observed; AR81009 and MS14003 uniquely induced 127 and 16 *G. hirsutum* genes, respectively (Fig 6c). In contrast, the average magnitude of gene induction between the two strains was not significantly different (Fig S4). Both *Xcm* strains caused more genes to be differentially expressed in DES 56 than in Acala Maxxa. Among the 52 genes significantly induced by both strains, sixteen conserved targets are homeologous pairs, whereas seventeen and fifteen genes are encoded by the A and D sub-genomes, respectively (Tables 2 and S4). It has been previously reported that homeologous genes encoded on the *G. hirsutum* A and D sub-genomes are differentially regulated during abiotic stress [46]. A set of approximately 10,000 homeologous gene pairs were selected and differential gene expression was assessed (Fig 7). For each pairwise comparison of *Xcm* strain and *G*. *hirsutum* cultivar, a similar number of genes were differentially expressed in each of the A and D subgenomes. However, some homeologous pairs were up- or down-regulated differentially in response to disease, indicating a level of sub-genome specific responses to disease. For example, SWEET sugar transporter gene Gh_D12G1898 in the D genome is induced over fourfold during infection with *Xcm* strain AR81009, while the homeolog Gh_A12G1747 in the A genome is induced to a much smaller extent.

**Figure 6.**
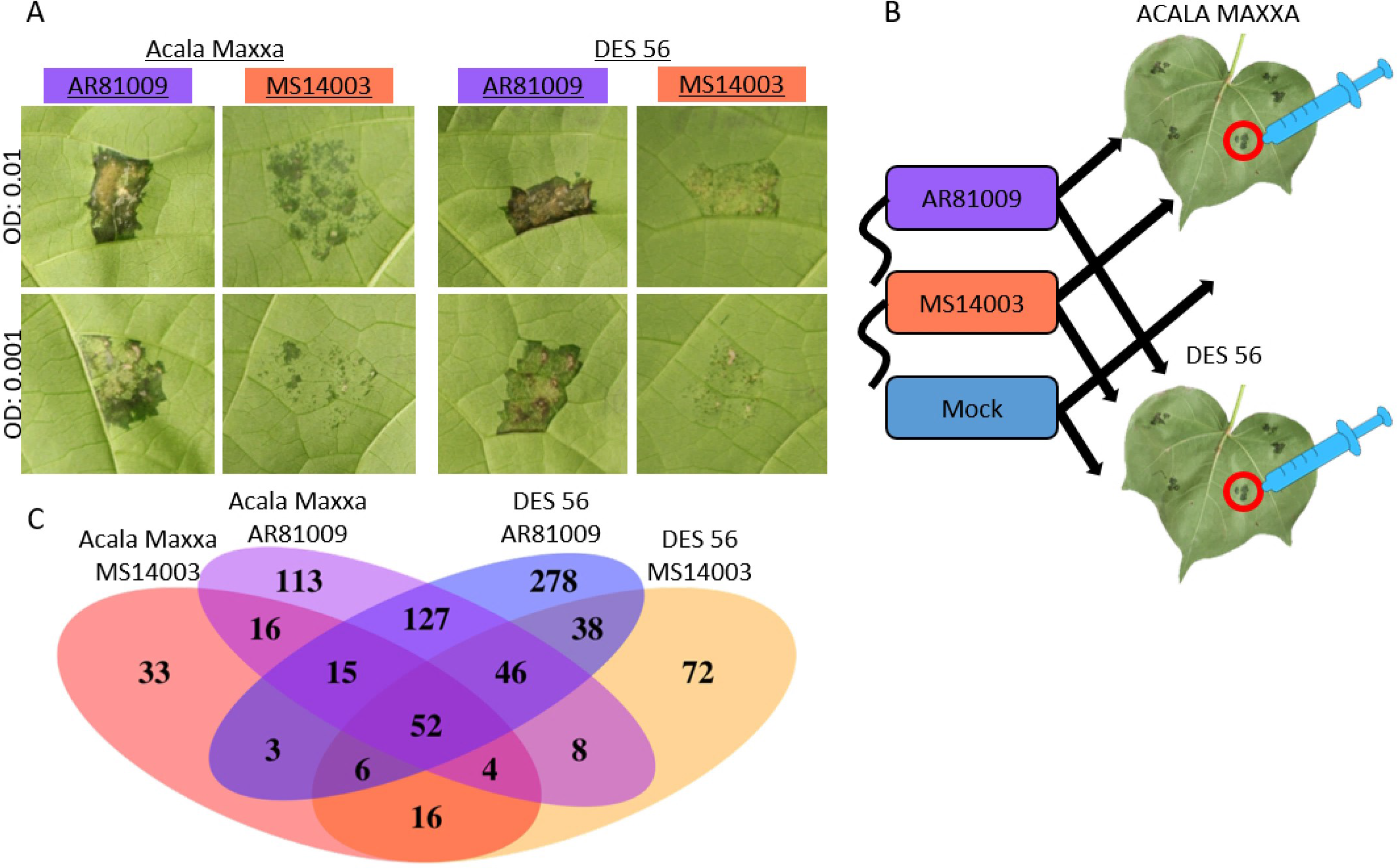
RNA-Sequencing analysis of infected *G. hirsutum* tissue demonstrates transcriptional changes during CBB. A) Disease phenotypes of *Xcm* strains MS14003 and AR81009 on *G. hirsutum* cultivars Acala Maxxa and DES 56, 7 days post inoculation. B) Acala Maxxa and DES 56 were inoculated with *Xcm* strains MS14003 and AR81009 at an OD of 0.5 and a mock treatment of 10mM MgCl_2_. Inoculated leaf tissue was collected at 24 and 48 hpi (before disease symptoms emerged). C) Venn diagram of upregulated *G. hirsutum* genes (Log2(fold change in FPKM) ≥ 2 and p value ≤ 0.05) in response to *Xcm* inoculation. Venn diagram was created using the VennDiagram package in R.

**Figure 7.**
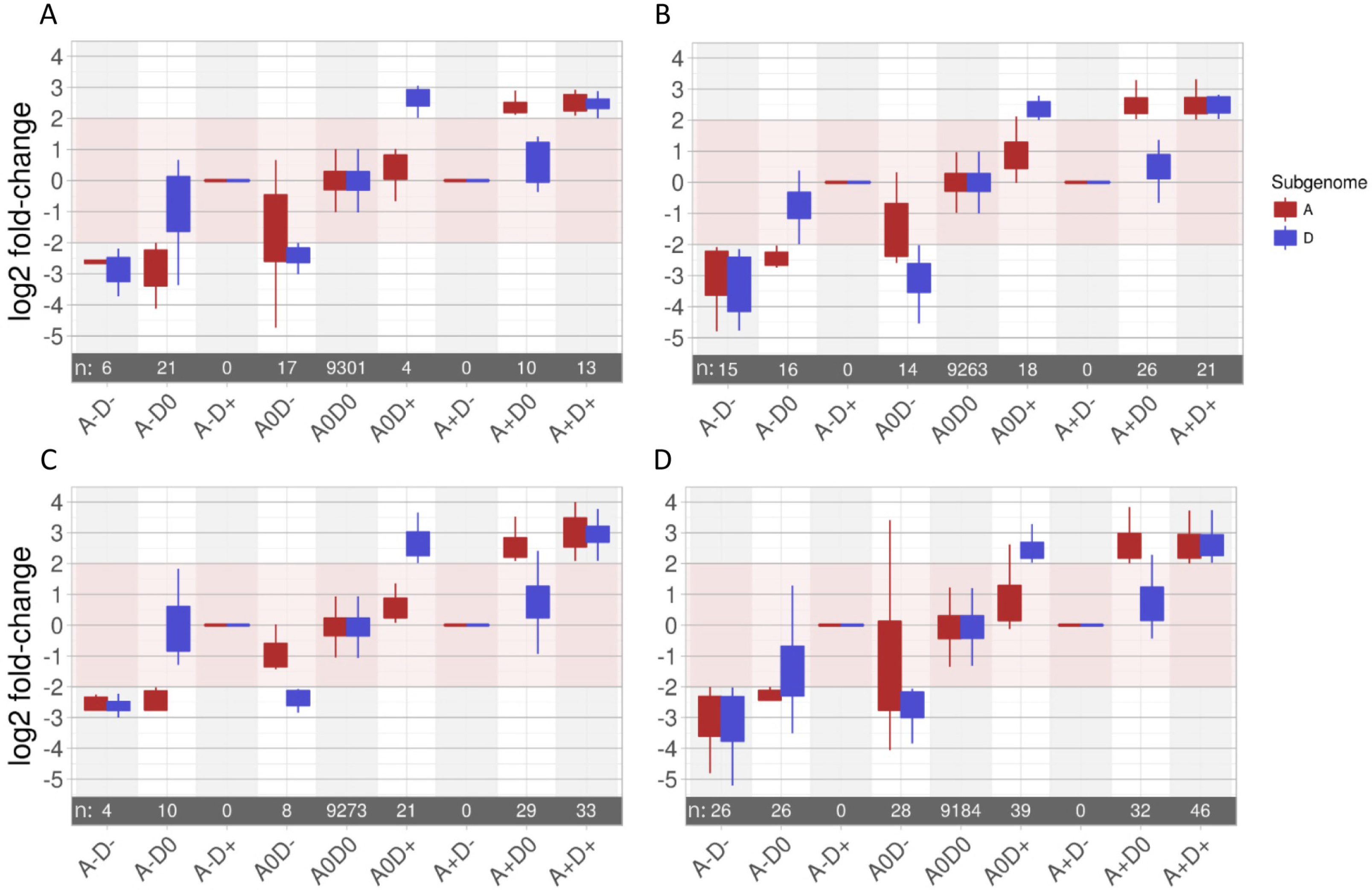
Expression of homeologous pairs across the A and D *G. hirsutum* genomes in response to *Xcm* inoculation. Genes are considered up or down regulated if the absolute value of gene expression change after inoculation as compared to mock treatment was Log2(fold change in FPKM) ≥ 2 and p value ≤ 0.05. By these criteria, pink shading indicates no significant gene expression change. A-D-: both members of the homeologous gene pair are down regulated; A- D0: only the ‘A’ sub-genome homeolog is down regulated; A-D+: ‘A’ sub-genome homeolog is down regulated, ‘D’ sub-genome homeolog is upregulated; etc. Number of gene pairs (n) meeting each expression pattern is indicated within the grey bar. For all genes meeting each expression pattern, the distribution of expression patterns is displayed as a box plot. Rectangles indicate the interquartile range and the whiskers show 1.5 times the interquartile range. A) Acala Maxxa inoculated with MS14003 B) DES 56 inoculated with MS14003 C) Acala Maxxa inoculated with AR 81009 D) DES 56 inoculated with AR81009.

**Table 2.**
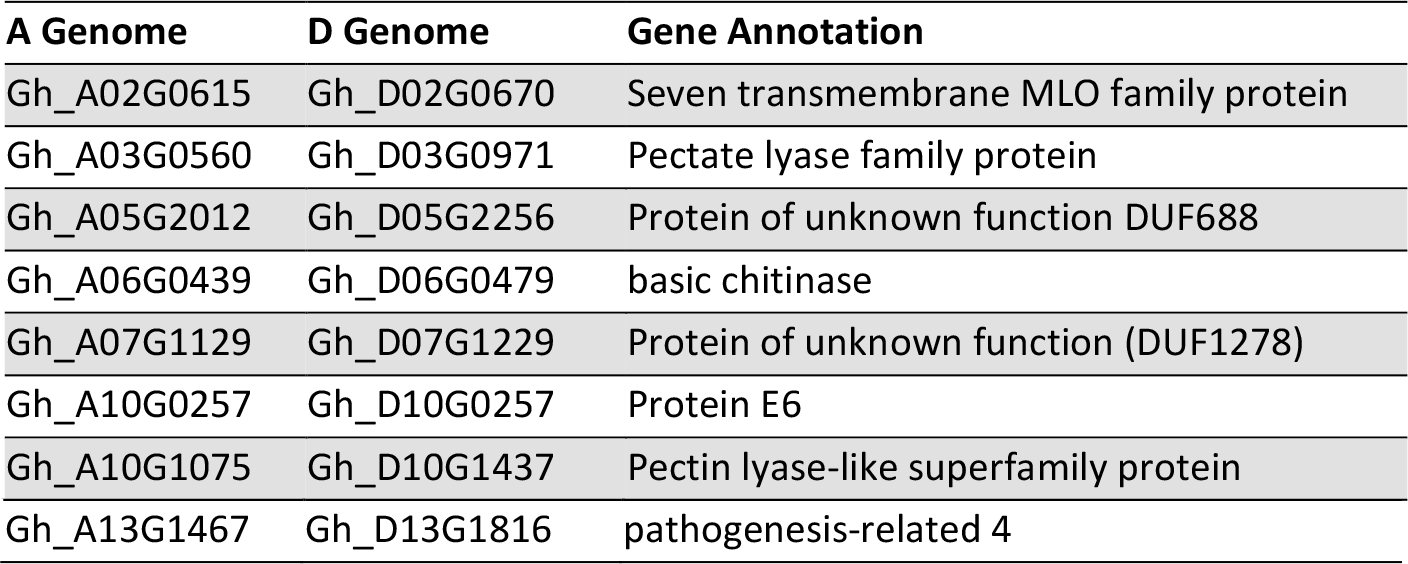
Eight Homeologous pairs of *Gossypium hirsutum* genes upregulated in Acala Maxxa and DES 56 varieties 48 hours post inoculation with *Xanthomonas citri pv. malvacearum* strains MS14003 and AR81009.

### Different strains of Xcm target distinct SWEET transporters in G. hirsutum

SWEET sugar transporter genes have been reported to be targets of and upregulated by *Xanthomonas* TAL effectors in *Manihot esculenta*, *Oryza sativa*, and *Citrus sinensis* [21, 40, 47, 48]. In rice and cassava, the SWEET genes are confirmed susceptibility genes that contribute to disease symptoms. The previously reported susceptibility genes and the SWEETs identified here, are clade III sugar transporters (Fig S5). The NBI *Gossypium hirsutum* genome encodes 54 putative SWEET sugar transporter genes. Of these 54 genes, three were upregulated greater than fourfold in response to inoculation by one of the two *Xcm* strains (Fig 8). Predicted TAL effector binding sites were identified using the program TALEnt [49]. MS14003 significantly induces the homeologs Gh_A04G0861 and Gh_D04G1360 and contains the TAL effectors M14b, M28a, and M28b, which are predicted to bind within the 300bp promoter sequences of at least one of these genes. Of note is TAL M28a, which is predicted to bind both homeologs (Fig S6a). In contrast, AR81009 induces Gh_D12G1898 to a greater extent than its homeolog Gh_A12G1747. TAL effectors A14c and A16b from AR81009 are predicted to bind to the Gh_D12G1898 and Gh_A12G1747 promoters; however, TAL A14a is predicted to bind only the Gh_D12G1898 promoter (Fig S6b). We note that while Gh_A12G1747 did not pass the fourfold cut off for gene induction, this gene is slightly induced compared to mock inoculation.

**Figure 8.**
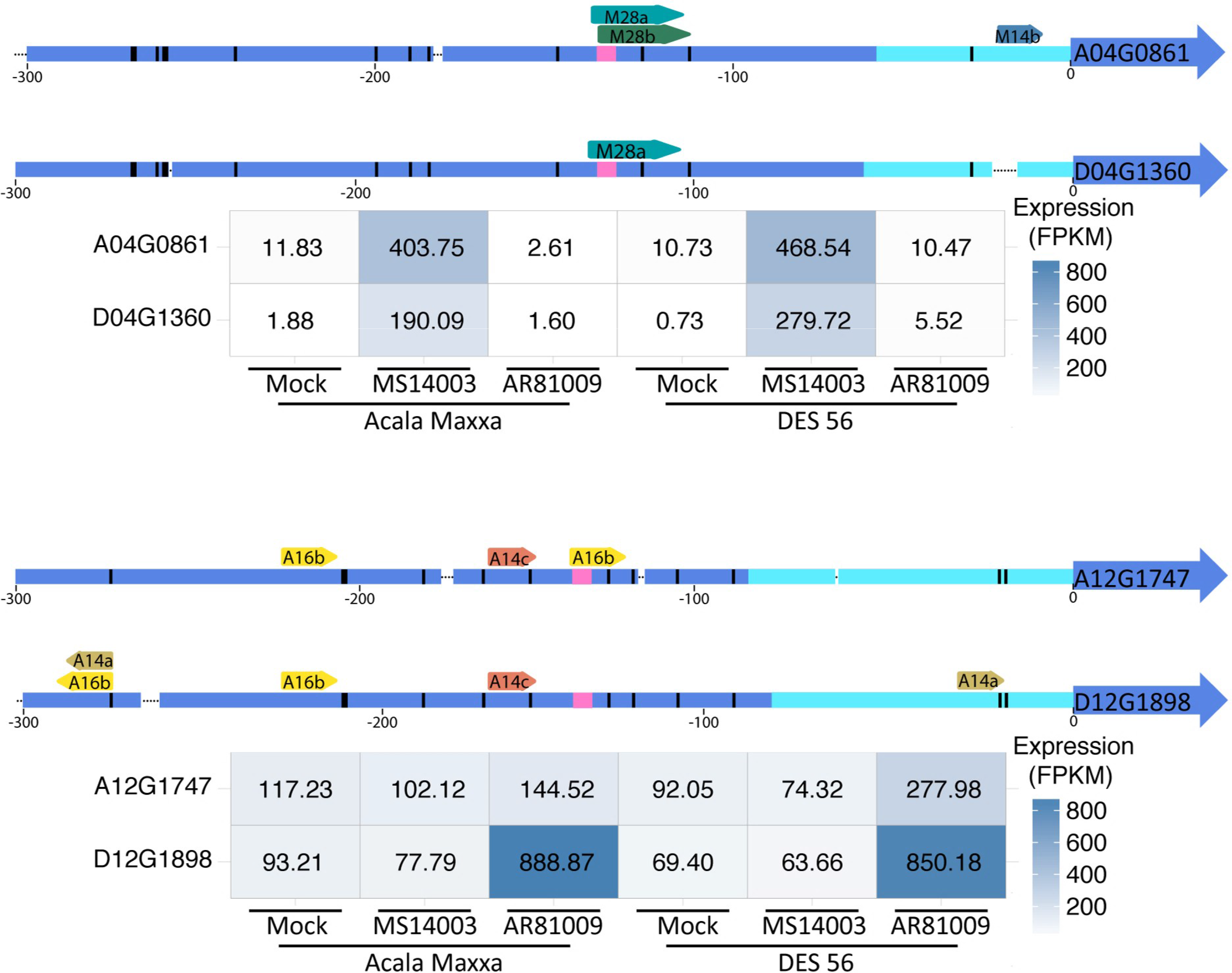
Three candidate *G. hirsutum* susceptibility genes are targeted by two different *Xcm* strains. A) The homeologous pair of SWEET genes A04_G0861 and D04_G1360 are upregulated in the presence of *Xcm* strain MS14003. (top) Cartoon summary of 300bp promoters of A04_G0861 and D04_G1360. (bottom) Heat-map of the expressions of A04_G0861 and D04_G1360 48 hours after mock or *Xcm* inoculation. B) The SWEET gene D12_G1898 is upregulated in the presence of *Xcm* strain AR81009. (top) Cartoon summary of 300bp promoters of D12_G1898 and A12_G1747. (bottom) Heat-map of the expressions of A12_G1747 and D12_G1898 48 hours after mock or *Xcm* inoculation. TAL effector binding sites were predicted with TALEsf using a quality score cutoff of 4. Gene promoter cartoon legend: Arrow: TAL effector binding site; Black dot: Deletion; Black bar: SNP; Pink bar: TATA box; Teal section: 5’UTR.

## Discussion

Cotton Bacterial Blight was considered controlled in the U.S. until an outbreak was observed during the 2011 growing season in Missouri, Mississippi and Arkansas [50]. Until 2011, seed sterilization, breeding for resistant varieties, and farming techniques such as crop rotation and sterilizing equipment prevented the disease from becoming an economic concern [51]. The number of counties reporting incidence of CBB has increased from 17 counties in 2011 to 77 counties in 2015 [38, 52, 53]. This paper investigates the root of the re-emergence and identifies several routes towards control of the disease.

When CBB was first recognized as re-emerging, several possible explanations were proposed including: (1) A highly virulent race of the pathogen that had been introduced to the U.S.; (2) Historical strains of *Xcm* that had evolved to overcome existing resistance (e.g. an effector gene change or host shift); and (3) Environmental conditions over the last several years that had been particularly conducive to the disease. Here, we present evidence that the re-emergence of CBB is not due to a large genetic change or race shift in the pathogen. Rather, the re-emergence of the disease is likely due to agricultural factors such as large areas of susceptible cultivars being planted. The presented data do not rule out potential environmental conditions that may also have contributed to the re-emergence. In this context, environmental conditions include disease conducive temperature and humidity as well as potentially contaminated seed or other agronomic practices that may have perpetuated spread of the disease outbreaks. Importantly, the presented data confirm that the presence of resistance loci could be deployed to prevent further spread of this disease. However, since many of the most popular farmer preferred varieties lack these resistance traits, additional breeding or biotechnology strategies will be needed to maximize utility. Notably, the current *Xcm* isolates characterized in this study all originate from Mississippi cotton fields in 2014. During the 2015 and 2016 growing seasons, resistant cotton cultivars were observed in Texas with symptoms indicative of bacterial infection distinct from CBB. Additional work is underway to identify and characterize the causal agent(s) of these disease symptoms.

Recent work on CBB in the US has focused on the most prevalent US *Xcm* race: race 18. However, races are not necessarily phylogenetically distinct clades. Race 18 isolates have been reported overseas, indicating that there may be independent origins of the race or cross-continent movement of this pathogen. Phenotypic race delineations were created before modern genetic and phylogenetic techniques were developed. However, modern genetics presents the opportunity to begin classifying strains based upon phylogenetic and effector profiles rather than phenotypes on a limited range of host varieties. Here, we identify all known and putative race 18 isolates as phylogenetically grouped into a single clade and distinct from other *Xcm* isolates. Future efforts can further explore phylogenetic relatedness among diverse isolates.

While resistant cotton cultivars were identified for all strains in this study, variability in symptom severity was observed for different strains when inoculated into susceptible cultivars. Two strains in particular, MS14003 and AR81009, have different effector profiles as well as different disease phenotypes. Comparative genomic analysis of the two pathogens revealed many differences that may contribute to the relative disease severity phenotypes. Similarly, transcriptomic analysis of two cultivars of *G. hirsutum* inoculated with these strains confirm that the genomic differences between the two strains result in a divergence in their molecular targets in the host.

Over the past decade, susceptibility genes have become targets for developing disease tolerant plants [54, 55]. These genes are typically highly induced during infection [56]. Therefore, RNA-Seq of infected plants has become a preferred way to identify candidate susceptibility genes. Once identified, genome editing can be used to block induction of these genes [57]. We report a homeologous pair of genes that are homologs of the MLO gene as targeted by both *Xcm* strains in both cotton cultivars. These genes are excellent candidates for future biotechnology efforts. Because the potential importance of these genes in cotton biology is unknown, their role in cotton physiology must first be explored. Knock-out mutations of MLO genes in other systems has led to durable resistance against powdery mildew as well as oomycetes and bacteria such as *Xanthomonas* [42, 45]. The dual purpose of host susceptibility genes has been observed previously. For example, the rice *Xa13* (aka. *Os8N3* and *OsSWEET11*) gene is required for pollen development but also targeted by a rice pathogen during infection [58]. Xa13 is a member of the clade III SWEET sugar transporters implicated in many pathosystems. In this case, the induction of *Xa13* for pathogen susceptibility is mediated by a TAL effector. Of the 54 SWEET genes in the *G. hirsutum* genome, at least three are significantly upregulated during *Xcm* infection. In contrast to MLO, no single SWEET gene was induced by both pathogen strains in both hosts.

Analysis of SWEET gene expression after inoculation revealed a context for polyploidy in the *G. hirsutum*-*Xcm* pathosystem. This relatively unexplored area of plant-microbe interactions arose from our observation of a potential difference in induction magnitude between the homeologous Gh_A12G1747 and Gh_D12G1898 SWEET genes. Further analysis revealed many examples of preferentially induced or down-regulated homeologs in response to *Xcm* infection. Characterization of sub-genome specialization may lead to new insights regarding durability of resistance and susceptibility loci in polyploid crops. Future research may investigate the diploid ancestors of tetraploid cotton to further explore the evolution of host and pathogen in the context of ploidy events [59]

Multiple putative TAL effector binding sites were identified within each up-regulated SWEET promoter. These observations suggest that TAL M28a from MS14003 may induce the homeologs Gh_A04G0861 and Gh_D04G1360. Further, TAL effector A14a from AR81009 is likely responsible for the upregulation of Gh_D12G1898. Whether additional TAL effectors are involved in these responses is not clear. Genome organization in the host, such as histone modifications or other epigenetic regulations may also be affecting these interactions. Future research will investigate these mechanisms further.

Collectively, the data presented here suggest that the wide-spread planting of CBB-susceptible cultivars has contributed to the re-emergence of CBB in the southern U.S. It is possible that a reservoir of race 18 *Xcm* was maintained in cotton fields below the level of detection due to resistant cultivars planted in the 1990s and early 2000s. Alternatively, the pathogen may have persisted on an alternate host or was re-introduced by contaminated seed [9, 10]. Regardless of the cause of the re-emergence, the genomic comparisons among pathogen races and host cultivars has identified several possible routes towards resistance. These include the use of existing effective resistance loci as well as the potential disruption of the induction of susceptibility genes through genome editing. The latter is an attractive strategy in part because of recent progress in genome editing [60, 61]. In summary, within a relatively short time frame, through the deployment of modern molecular and genomic techniques, we were able to identify factors that likely contribute to the re-emergence of cotton bacterial blight and generate data that can now be rapidly translated to effective disease control strategies.

## Materials & Methods

### Xcm strain isolation and manipulation

New *Xcm* strains were isolated from infected cotton leaves by grinding tissue in 10mM MgCl_2_ and culturing bacteria on NYGA media. The most abundant colony type was selected, single colony purified and then 16S sequencing was used to confirm the bacterial genus as previously described [62]. In addition, single colony purified strains were re-inoculated into cotton leaves and the appearance of water soaked symptoms indicative of CBB infection was confirmed. Both newly isolated strains as well as strains received from collaborators were used to generate a rifampicin resistance version of each strain. Wildtype strains were grown on NYGA, then transferred to NYGA containing 100µg/ml rifampicin. After approximately 4-5 days, single colonies emerged. These were single colony purified and stored at -80C. The rifampicin resistant version of each *Xcm* strain was used in all subsequent experiments reported in this manuscript unless otherwise noted.

### Plant inoculations

Cotton varieties from the original cotton panel for determining *Xcm* race designations were obtained from the USDA/ARS, Germplasm Resources Information Network (GRIN). Varieties included in the *G. hirsutum* NAM population were provided by Vasu Kuraparthy [33]. Other commercial varieties were obtained from Terry Wheeler and Tom Allen. Disease assays were conducted in a growth chamber set at 30°C and 80% humidity. *Xcm* strains were grown on NYGA plates containing 100µg/ml rifampicin at 30°C for two days before inoculations were performed. Inoculations were conducted by infiltrating a fully expanded leaf with a bacterial solution in 10mM MgCl_2_ (OD_600_ specified within each assay).

The field tests were conducted as follows: Cotton cultivars are planted in two row plots (10 – 11 m in length, 1 m row spacing), in a randomized complete block design with four replications. Approximately 60 to 80 days after planting, *Xcm* was applied to the test area similar to that described in Wheeler et al. (2007) [37]. Briefly, Xcm is grown in trypticase soy broth (30 g/L) for 1 ½ days and then 19 L of the concentrated bacterial solution (10^8^ cfu/ml) are diluted into 189 L of water (resulting in 10^6^ cfu/ml). The surfactant Silwet L-77 (polyalkyleneoxide modified heptamethyltrisiloxane, Loveland Industries, Greely, CO) is added at 0.2% v/v. The suspension of bacteria are sprayed over the top of the cotton at a pressure of 83 kpa and rate of 470 L/ha. The nozzles used were TeeJet 8008. Symptoms were typically visible 14 days after application and plots were rated for incidence of symptoms 17-21 days after application [34-37].

### Cotton Cultivar Statistics

Area of cotton planted per county in the United States in 2015 was obtained from the USDA National Agricultural Statistics Service: www.nass.usda.gov/Statistics_by_Subject/result.php?7061F36A-A4C6-3C65-BD7F-129B702CFBA2&sector=CROPS&group=FIELD%20CROPS&comm=COTTONUSDA. Estimated percentage of upland cotton planted for each variety was obtained from the Agricultural Marketing Service (AMS): www.ams.usda.gov/mnreports/canvar.pdf.

### Bacterial Sequencing and Phylogenetics

Illumina based genomic datasets were generated as previously described [29]. Paired-end Illumina reads were trimmed using Trimmomatic v0.32 (ILLUMINACLIP:TruSeq3-PE.fa:2:30:10 LEADING:3 TRAILING:3 SLIDINGWINDOW:4:15 MINLEN:36) [63]. Genome assemblies were generated using the SPAdes *de novo* genome assembler [64]. Strain information is reported in Supplemental Table 1. Similar to our previously published methods [29], the program Prokka was used in conjunction with a T3E database to identify type three effector repertoires for each of the 12 *Xcm* isolates as well as four *Xcm* genomes previously deposited on NCBI (S2Table) [65].

Multi-locus sequence analysis was conducted by concatenating sequences of the *gltA*, *lepA*, *lacF*, *gyrB*, *fusA* and *gap-1* loci obtained from the Plant-Associated Microbes Database (PAMDB) for each strain as previously described [66]. A maximum-likelihood tree using these concatenated sequences was generated using CLC Genomics 7.5.

### Variant Based Phylogeny

A variant based dendrogram was created by comparing 12 Illumina sequenced *Xcm* genomes to the complete *Xanthomonas citri* subsp. *citri* strain Aw12879 reference genome (Genbank assembly accession: GCA_000349225.1) on NCBI. Read pairs were aligned to the reference genome using Bowtie2 v2.2.9 with default alignment parameters [27]. From these alignments, single nucleotide polymorphisms (SNPs) were identified using samtools mpileup v1.3 and the bcftools call v1.3.1 multi-allelic caller [28]. Using Python v2.7, the output from samtools mpileup was used to identify loci in the *X. citri* subsp. *citri* reference genome with a minimum coverage of 10 reads in each *Xcm* genome used Python version 2.7 available at http://www.python.org. Vcftools v0.1.14 and bedtools v2.25.0 were used in combination to remove sites marked as insertions or deletions, low quality, or heterozygous in any of the genomes [67, 68]. Remaining loci were concatenated to create a FASTA alignment of confident loci. Reference loci were used where SNP's were not detected in a genome. The resulting FASTA alignment contained 17853 loci per strain. This alignment was loaded into the online Simple Phylogeny Tool from the ClustalW2 package to create a neighbor joining tree of the assessed strains [69, 70]. Trees were visualized using FigTree v1.4.2.

### Genome Assembly

Single Molecule, Real Time (SMRT) sequencing of *Xcm* strains MS14003 and AR81009 was obtained from DNA prepped using a standard CTAB DNA preparation. Blue Pippin size selection and library preparation was done at the University of Deleware Sequencin Facility. The genomes were assembled using FALCON-Integrate (https://github.com/PacificBiosciences/FALCON-integrate/commit/cd9e93) [71]. The following parameters were used: Assembly parameters for MS14003: length_cutoff = 7000; length_cutoff_pr = 7000; pa_HPCdaligner_option = -v -dal8 -t16 -e.70 -l2000 -s240 -M10; ovlp_HPCdaligner_option = -v -dal8 -t32 -h60 -e.96 -l2000 -s240 -M10; falcon_sense_option = – output_multi –min_idt 0.70 –min_cov 5 –local_match_count_threshold 2 –max_n_read 300 – n_core 6; overlap_filtering_setting = –max_diff 80 –max_cov 160 –min_cov 5 –bestn 10; Assembly parameters for AR81009: length_cutoff = 8000; length_cutoff_pr = 8000; pa_HPCdaligner_option = -v -dal8 -t16 -e.72 -l2000 -s240 -M10; ovlp_HPCdaligner_option = -v - dal8 -t32 -h60 -e.96 -l2000 -s240 -M10; falcon_sense_option = –output_multi –min_idt 0.72 – min_cov 4 –local_match_count_threshold 2 –max_n_read 320 –n_core 6; overlap_filtering_setting = –max_diff 90 –max_cov 300 –min_cov 10 –bestn 10. Assemblies were polished using iterations of pbalign and quiver, which can be found at https://github.com/PacificBiosciences/pbalign/commit/cda7abb and https://github.com/PacificBiosciences/GenomicConsensus/commit/43775fa. Two iterations were run for *Xcm* strain MS14003 and 3 iterations for AR81009. Chromosomes were then reoriented to the DnaA gene and plasmids were reoriented to ParA. The assemblies were checked for overlap using BLAST, and trimmed to circularize the sequences [72]. TAL effectors were annotated and grouped by RVD sequences using AnnoTALE [41]. Homologous regions among plasmids that are greater than 1 kb were determined using progressiveMauve [39]. Genomic comparisons between the MS14003 and AR81009 chromosomes were visualized using Circos [73]. Single-copy genes on each of the chromosomes were identified and joined using their annotated gene IDs. Lines connecting the two chromosomes represent these common genes and their respective positions in each genome. A sliding window of 1KB was used to determine the average GC content. Methylation was determined using the Base Modification and Motif Analysis workflow from pbsmrtpipe v0.42.0 at https://github.com/PacificBiosciences/pbsmrtpipe.

### Western Blot Analysis

Western Blot analysis of Transcription Activator-Like (TAL) effectors was performed using a polyclonal TAL specific antibody [40]. Briefly, bacteria were suspended in 5.4 pH minimal media for 4.5 hours to induce effector production and secretion. Bacteria were pelleted and then suspended in laemmli buffer and incubated at 95 degrees Celsius for three minutes to lyse the cells. Freshly boiled samples were loaded onto a 4-6% gradient gel and run for several hours to ensure sufficient separation of the different sized TAL effectors.

### Gene Expression Analysis

Susceptible cotton were inoculated with *Xcm* using a needleless syringe at an OD_600_ of 0.5. Infected and mock-treated tissue were collected and flash frozen at 24 and 48 hours post inoculation. RNA was extracted using the Sigma tRNA kit. RNA-sequencing libraries were generated as previously described [74].

Raw reads were trimmed using Trimmomatic [63]. The Tuxedo Suite was used for mapping reads to the TM-1 NBI *Gossypium hirsutum* genome [75], assembling transcripts, and quantifying differential expression [27].

Read mapping identified several mis-annotated SWEET genes that skewed differential expression results. The annotations of SWEET genes Gh_A12G1747, Gh_D07G0487, and Gh_D12G1898 were shortened to exclude 20-30kb introns. Two exons were added to Gh_D05G1488. The 2.7kb scaffold named Scaffold013374 was also removed from analysis because its gene Gh_Sca013374G01 has exact sequence homology to Gh_A12G1747 and created multi-mapped reads that interfered with expression analysis.

Homeologous pairs were identified based on syntenic regions with MCScan [76]. A syntenic region was defined as a region with a minimum of five genes with an average intergenic distance of two and within extended distance of 40. All other values were set to the default. Comparisons between homeologs was performed by examining cuffdiff differential expression and classifying them according to the sub-genome expression pattern. Genes considered up or down regulated meet both differential expression from mock significance of q-value < 0.05 and the absolute value of the log2 fold change is greater than 2.

### TAL Binding Sites

Bioinformatic prediction of TAL effector binding sites on the *G. hirsutum* promoterome was performed using the TAL Effector-Nucleotide Targeter (TALEnt) [50]. In short, the regions of the genome that were within 300 basepairs of annotated genes were queried with the RVD’s of MS14003 and AR81009 using a cutoff score of 4. Promiscuously binding TALs 16 from MS14003 and 16a from AR81009 were removed from analysis.

## Acknowledgements

The authors would like to acknowledge Dr. Robert Nichols for useful discussions throughout the presented research and preparation of this manuscript.

## Supporting Information Legends

**S1 Table:** US Counties with reported CBB incidence from 2009 to 2016.

**S2 Table:** Xanthomonas genomes previously deposited on NCBI that are referenced in this paper.

**S3 Table:** Disease phenotypes and percent acreage of commercial G. hirsutum varieties planted in the US from 2009-2016.

**S4 Table:** RNA-Seq analysis reveals that 52 genes are induced in all Xcm-G. hirsutum interactions at 48 hours ((p ≤ 0.05) with a Log2 (fold change in FPKM) ≥ 2).

**S1 Fig:**
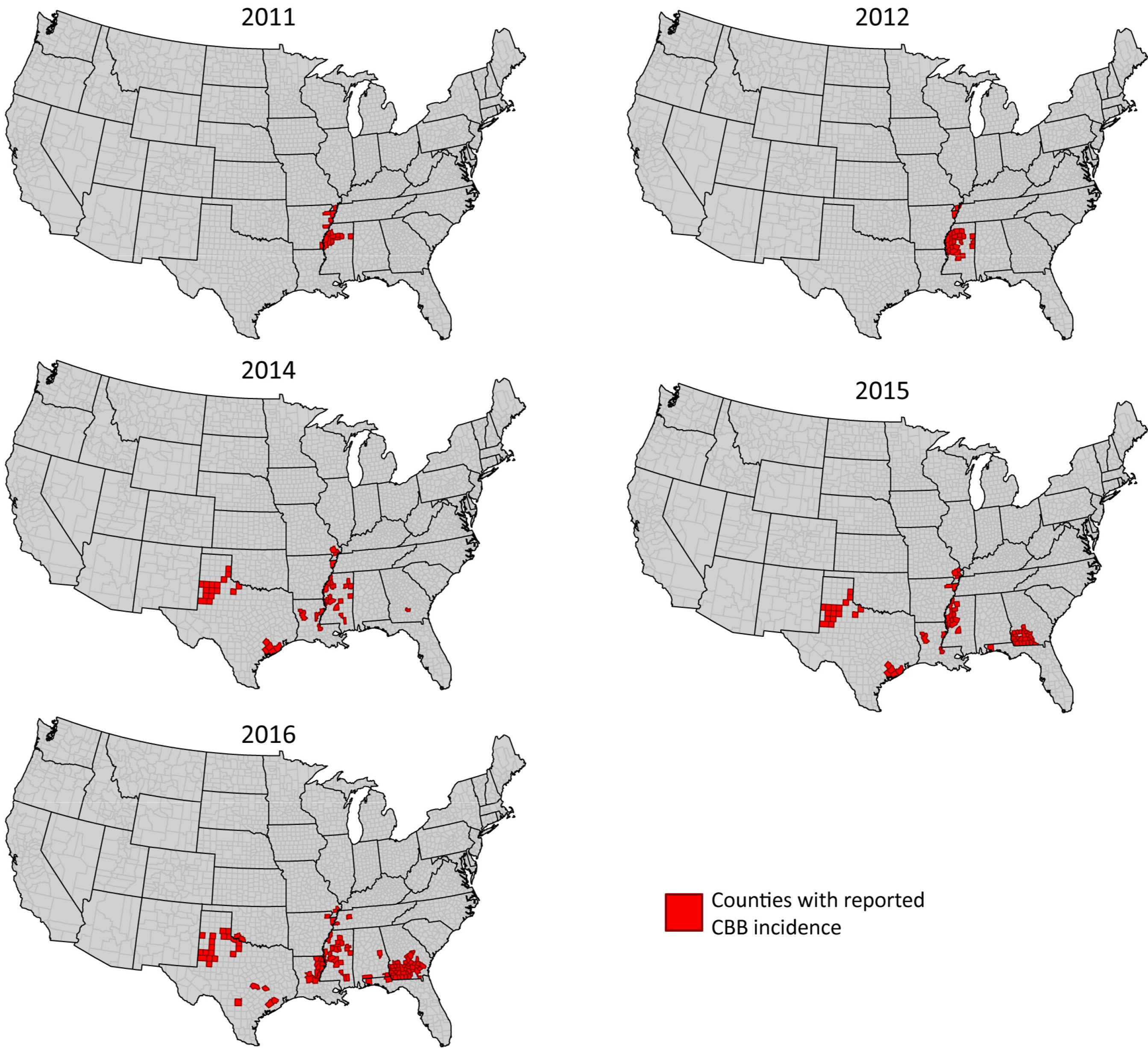
Maps of CBB incidence in the US from 2011-2012 and 2014-2016. CBB incidence was reported by extension agents, extension specialists and certified crop advisers in their respective states for the years 2011-2012 and 2014-2016, and compiled by Tom Allen. CBB reports for 2013 were infrequent.

**S2 Fig:**
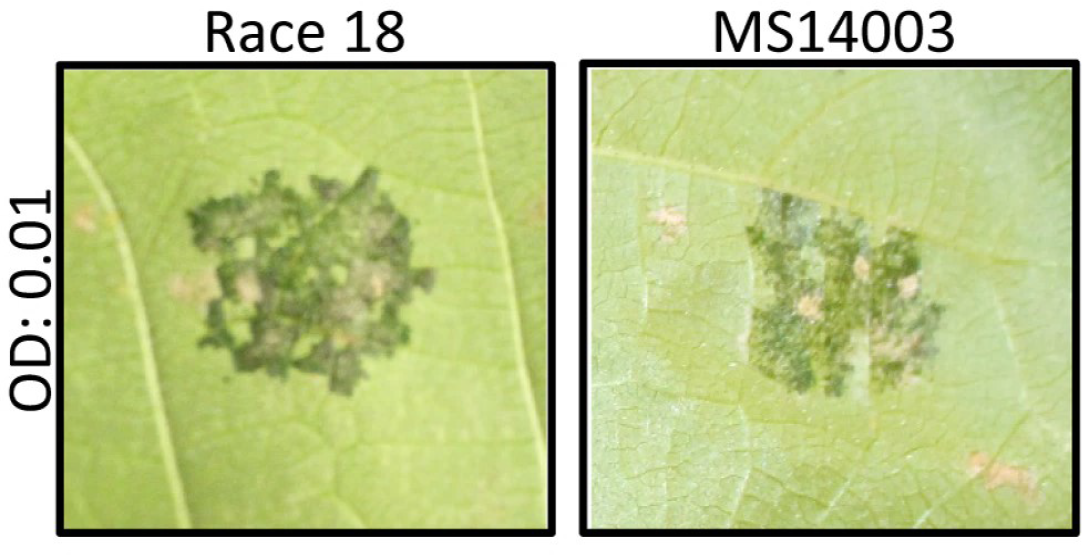
Disease phenotypes of historical Race18 strain and MS14003 strain. *Xcm* strains Race18 and MS14003 were inoculated into *G. hirsutum* variety PHY499 WRF at an OD600 of 0.01 and imaged at 8 dpi.

**S3 Fig:**
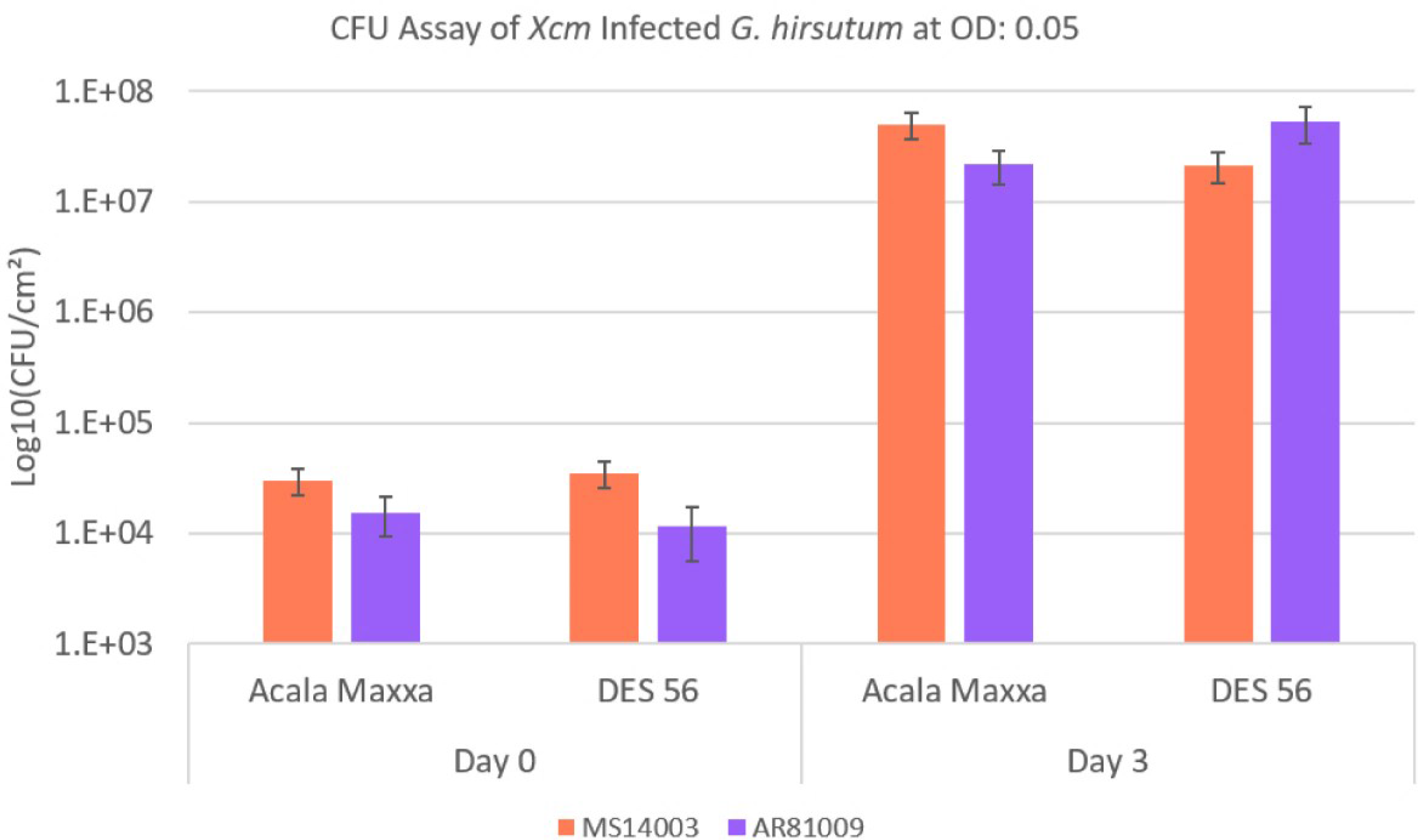
Growth assay of MS14003 and AR81009 on cotton varieties Acala Maxxa and DES 56. *G. hirsutum* varieties were inoculated with *Xcm* at an OD600: 0.05. Tissue was collected at day 0 and day 3 and processed as described in materials and methods.

**S4 Fig:**
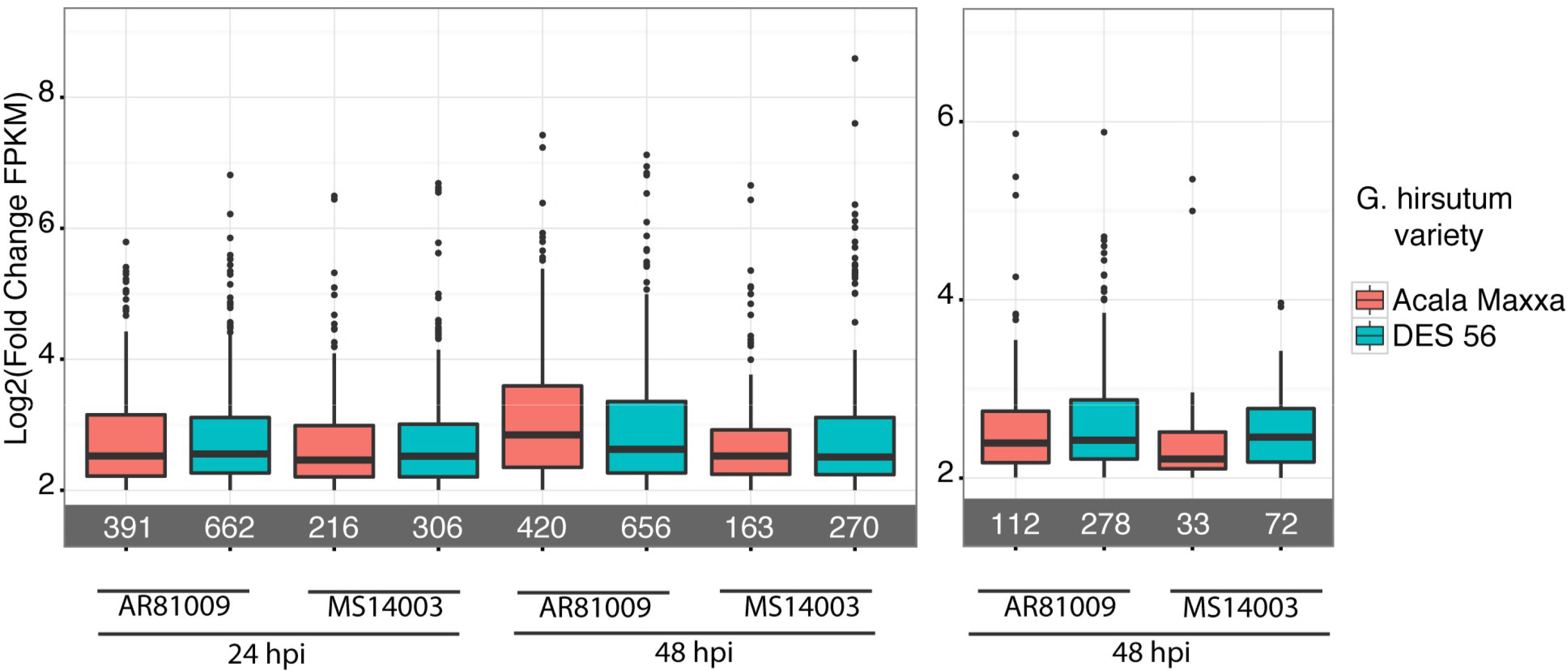
Expression levels of significantly upregulated genes with a Log2 fold change of 2 in *G. hirsutum*. A) All significantly upregulated genes with a Log2 fold change of 2 B) All significantly upregulated genes (p ≤ 0.05) with a Log2 (fold change in FPKM) ≥ 2 that are unique to each cultivar/*Xcm* disease interaction in *G. hirsutum*. Numbers in grey bar indicate the total number of genes for each condition.

**S5 Fig:**
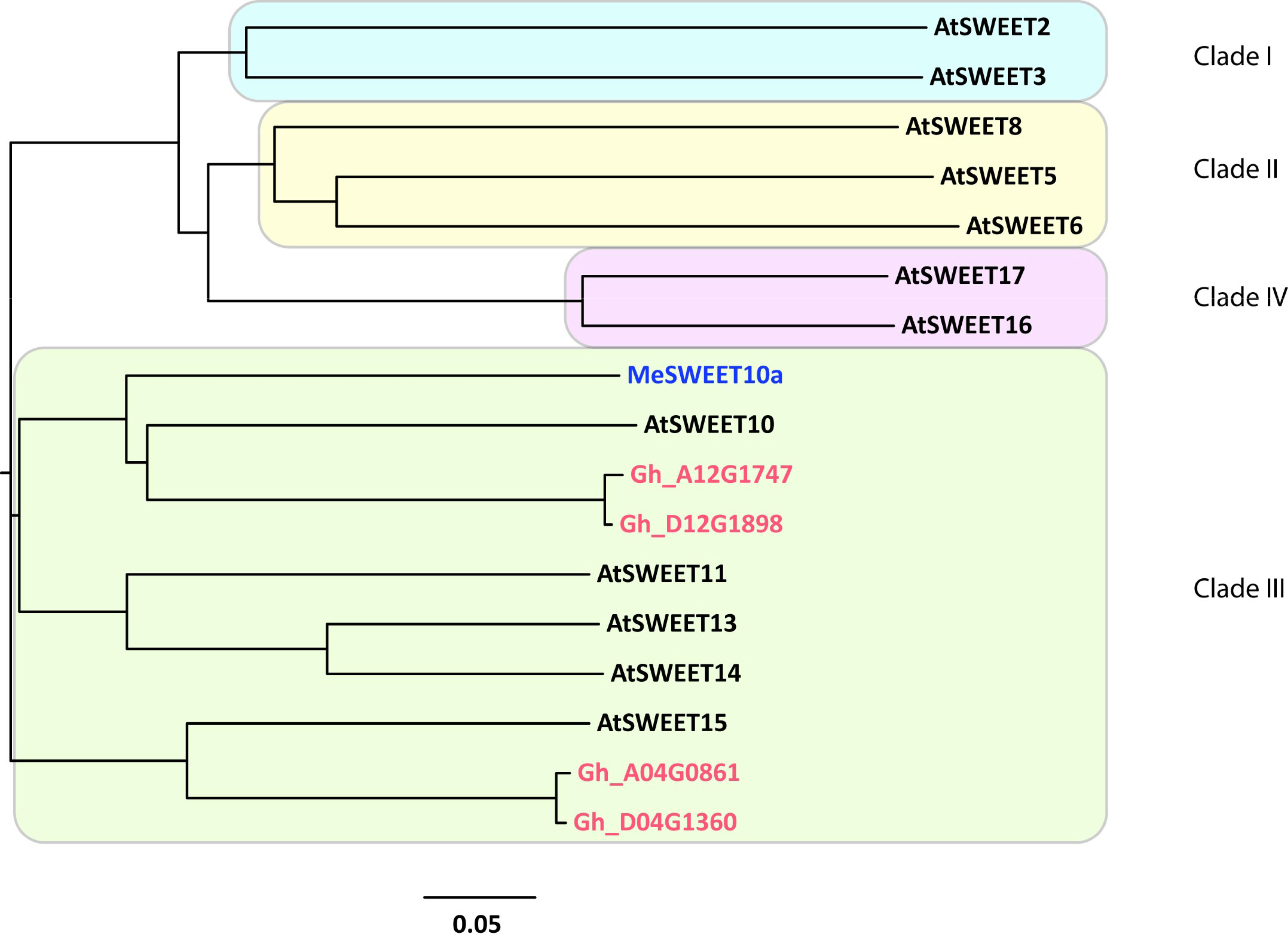
Phylogeny of SWEET genes from *Gossypium hirsutum*, *Manihot esculenta*, and *Arabidopsis thaliana*. Four predicted *G. hirsutum* SWEET genes are compared to classified *A. thaliana* SWEET genes and the MeSWEET10a *M. esculenta* susceptibility gene. A protein alignment and phylogenetic tree was generated by Clustal Omega, and the tree was visualized using Figtree v1.4.2.

**S6 Fig:**
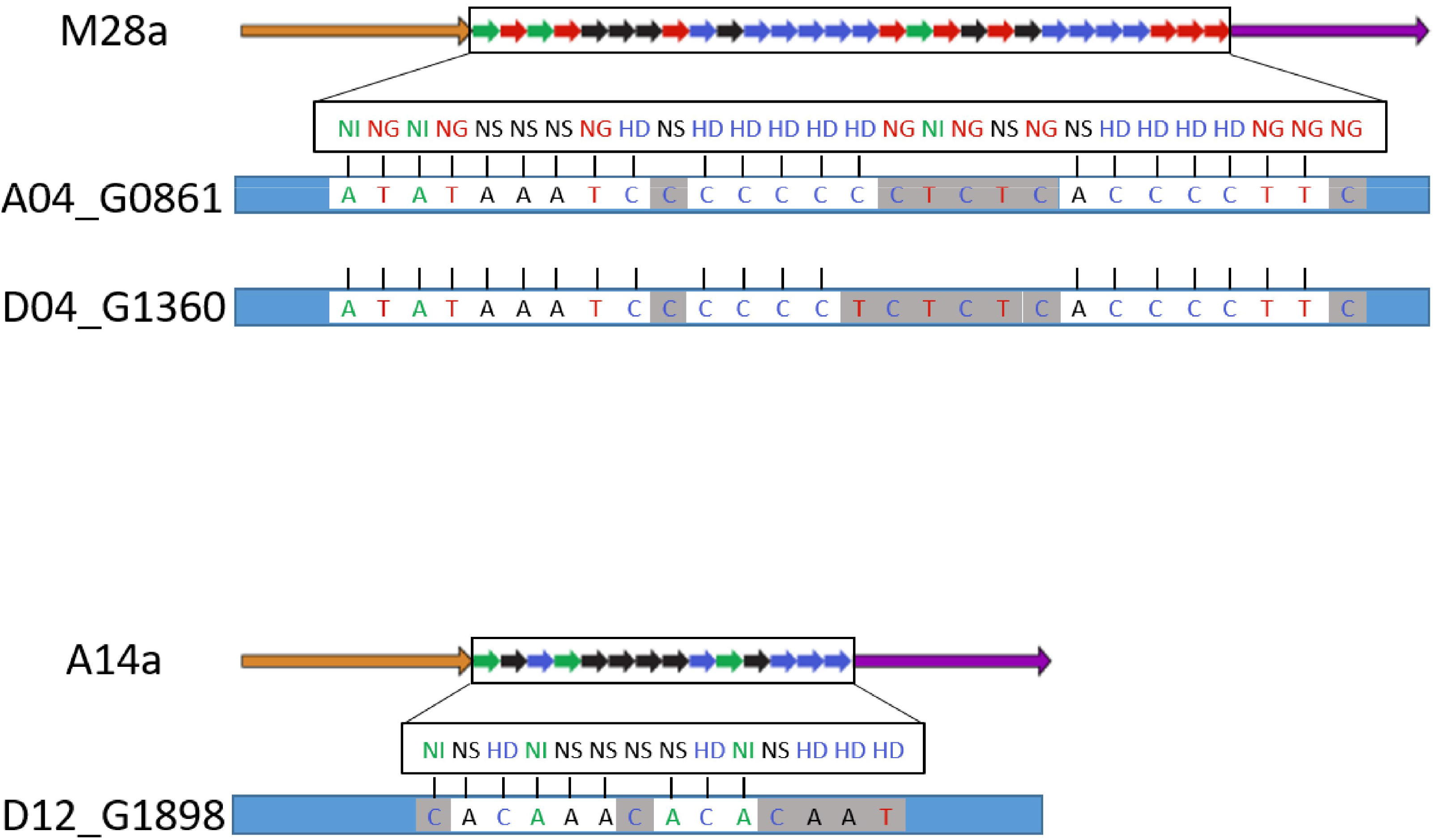
Alignment of predicted TAL effector binding sites on induced *G. hirsutum* SWEET genes. A) TAL M28a is predicted to bind to and up-regulate the homeologous pair of SWEET genes: A04_G0861 and D04_G1360 in *G. hirsutum* varieties Acala Maxxa and DES56 after inoculation with *Xcm* strain MS14003. B) TAL A14a is predicted to bind to and up-regulate the SWEET gene D12_G1898 G1360 in *G. hirsutum* varieties Acala Maxxa and DES56 after inoculation with *Xcm* strain AR81009.

